# *Anopheles albimanus* is a potential alphavirus vector in the Americas

**DOI:** 10.1101/2022.06.09.495519

**Authors:** Gerard Terradas, Mario Novelo, Hillery C. Metz, Marco Brustolin, Jason L. Rasgon

## Abstract

Despite its ecological flexibility and geographical co-occurrence with human pathogens, little is known about the ability for *Anopheles albimanus* to transmit arboviruses. To address this gap, we challenged *An. albimanus* females with four alphaviruses and one flavivirus and monitored progression of infections. We found this species is an efficient vector of the alphaviruses Mayaro virus, O’nyong-nyong virus, and Sindbis virus, though the latter two do not currently exist in its habitat range. *An. albimanus* was able to become infected with Chikungunya virus, but virus dissemination was rare (indicating the presence of a midgut escape barrier), and no mosquito transmitted. Mayaro virus rapidly established disseminated infections in *An. albimanus* females and was detected in the saliva of a substantial proportion of infected mosquitoes. Consistent with previous work in other anophelines, we find that *An. albimanus* is refractory to infection with flaviviruses, a phenotype that did not depend on midgut-specific barriers. Our work demonstrates that *An. albimanus* may be a vector of neglected emerging human pathogens and adds to recent evidence that anophelines are competent vectors for diverse arboviruses.

## Introduction

More than 400 species have been described for the genus *Anopheles,* with approximately 40 regarded as disease vectors of interest. Anopheline mosquitoes are primarily known for transmitting malaria^1^, but they also have the potential to transmit viruses^2,3^. In general, they are highly mobile and thrive by using human activities and movement to disperse around the globe^4,5^. While the most known and well-studied species are the African *An. gambiae* and the Indo-Iranian *An. stephensi* due to the malaria-associated socioeconomic and health burdens they cause in those regions^1^, less popular anopheline species predominate in other areas of the planet with the potential to spread different pathogens.

*Anopheles albimanus* is the main anopheline inhabiting northern South America, Central America and the Caribbean islands^6^. Its broad geographical distribution may be aided by the species’ ability to survive in both fresh^7,8^ and brackish water^9,10^. Though the species remains incompletely understood, *An. albimanus* has been described as a zoophilic, crepuscular, and exophagic mosquito^11,12,13^. However, host availability and environmental conditions appear to influence its host choice^14,15^ and resting behavior^16^. The flexible behavior of this species may be facilitating its spread into the southern USA, deeper into South America, and into urban areas where it encounters human hosts. Its expansion may also be helped by climate change, which is broadening the species’ geographical habitat range^6^. Despite the work that has been done to characterize its ecology and behavior, little is known about the capacity of the species to harbor and transmit classic and emerging tropical mosquito-borne viruses.

Many arboviruses produce similar disease symptoms in humans that include fever, headache, rash, diarrhea, and joint pain, which can last for months. Since treatment is not specific to the etiological agent and neither are many surveillance and diagnostic tools, the prevalence of emerging viruses can be misdiagnosed and hence underestimated in areas with more common viral outbreaks such as Chikungunya or dengue viruses, as initially occurred with the Zika virus epidemic^17^. However, despite displaying similar clinical symptoms, each virus interacts with the mosquito differently to achieve a successful human-to-mosquito-to-human viral transmission route. Arboviruses rely on the rapid infection of a mosquito after feeding on an infectious host, and must penetrate and overcome multiple tissue and immune barriers to propagate throughout the body and reach the salivary glands^18^. The virus must also replicate in the salivary glands efficiently to later infect a naive vertebrate host through salivation during a second bloodmeal. To date, only one arbovirus is known to be primarily transmitted through the bite of *Anopheles* mosquitoes in the field (O’nyong-nyong virus^19^, *An. funestus* and *An. gambiae),* but other *Anopheles* species have been shown to be capable vectors of alphaviruses in the lab^20,21^.

Here we ask whether An. *albimanus* is a competent vector of arboviruses. We orally challenged adult females with infectious bloodmeals containing one of the following togaviruses (genus *Alphavirus):* Mayaro (MAYV; -D and -L genotypes), Chikungunya (CHIKV), O’nyong-nyong (ONNV) and Sindbis virus (SINV), as well as the flavivirus dengue virus serotype 2 (DENV2), due to its prevalence and health impact worldwide. Collectively, these togaviruses produce human disease that spans the planet (Figure 1a): MAYV in Central/South America, ONNV in Africa, CHIKV across the tropics, and SINV in colder climates. The tested viruses have differences in both virion structure and envelope proteins that create variation in their capacity for cellular entry and invasion (Figure 1b). Following viral challenge, we monitored mosquitoes for arbovirus infection, dissemination of virus beyond the midgut and through the body, and secretion of virus in saliva. We report for the first time that *An. albimanus* can become infected with and transmit multiple alphaviruses—including MAYV, a human pathogen that is already spreading within this mosquito’s geographic range in the Americas.

**Figure 1.**
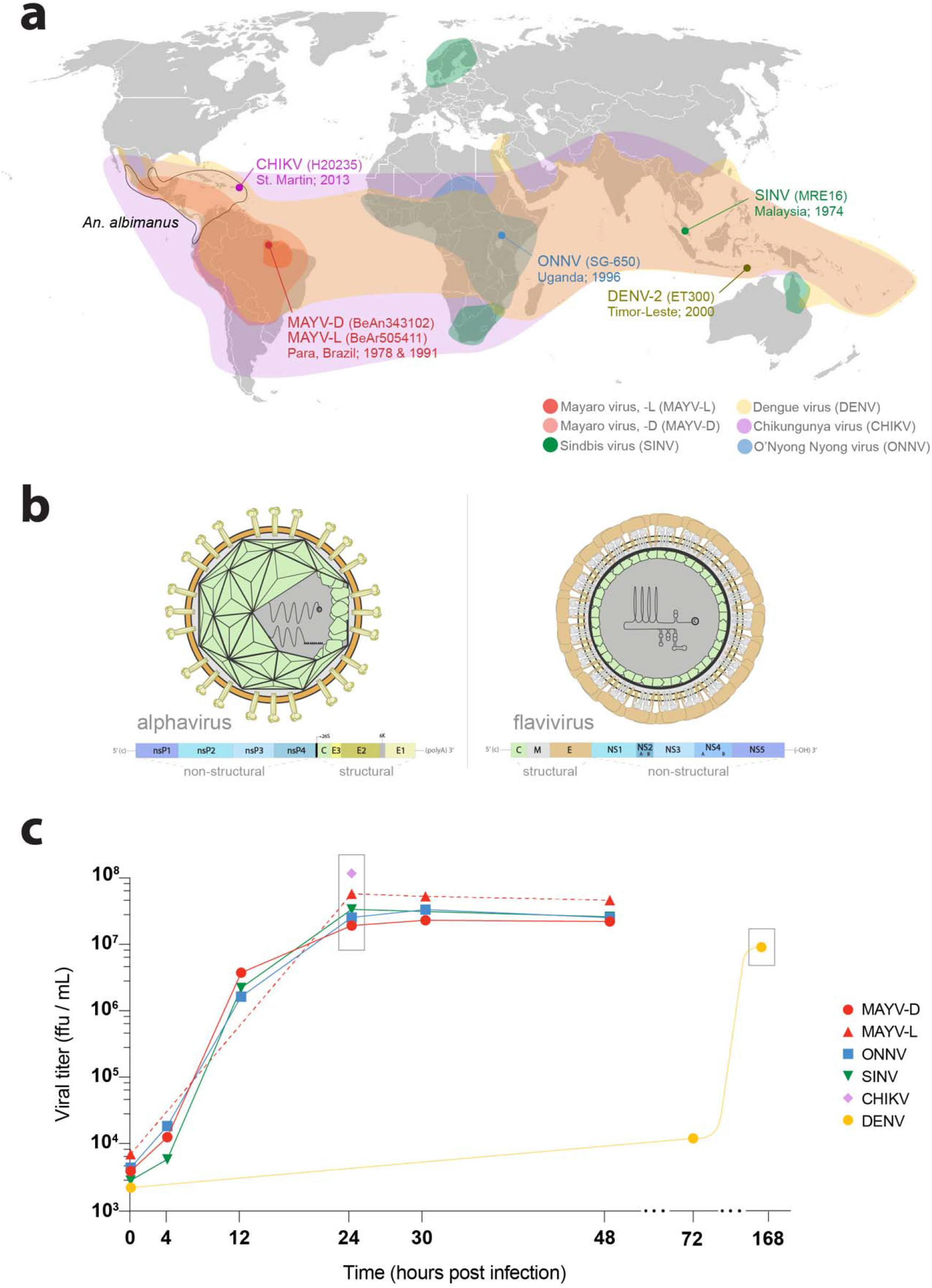
**a)** Geographical distribution of the arboviruses assessed in this study mapped with the geographical range of the Caribbean mosquito vector *An. albimanus* (black), **b)** Virion and genomic structure of alphaviruses (MAYV, ONNV, CHIKV, SINV) and flaviviruses (DENV), c) Alphavirus (up to 48 h) and flavivirus (up to 7 days) stocks were grown using Vero or C6/36 cells, respectively. Growth curves depict increasing infectious viral loads over time, from inoculation to final collection. The titers used for mosquito challenge experiments are shown within grey boxes.

## Results

### Growth kinetics of arboviruses *in vitro*

Prior to *in vivo* infection experiments, we assessed the *in vitro* growth of each virus over time. Each virus was propagated in Vero (alphaviruses) or C6/36 (flavivirus) cells. Alphaviruses typically replicate more quickly than flaviviruses, which can be seen in the rapid and severe cytopathic effects they produce in invaded cells of vertebrate origin^22^. We therefore collected samples for alphaviruses across a 48h span, when presence of abundant cellular death and abnormal media pH were visibly evident. Infectious titers of each virus stock for each timepoint were assessed by plaque forming assay (PFA, SINV), or focus forming assay (FFA, the remaining viruses). All alphavirus titers peaked at 24h post infection with viral titers >10^7^ ffu (or pfu)/mL (Figure 1c), after which they declined only slightly until the final timepoint (48h). Note that CHIKV’s viral titer (3×10^8^ ffu/mL) was assessed only at 24h post infection due to BSL-3 laboratory constraints. Conversely, DENV2 infections were sampled at day 0, 3 and 7 and we found viral titer in supernatant was highest at day 7 (1×10^7^ ffu/mL).

### Both MAYV genotypes infect, replicate, and are transmitted efficiently by *An. albimanus*

Our study focused on the potential ability of An. *albimanus* to transmit arboviruses, including those that are endemic within its geographic range such as MAYV, CHIKV, and DENV2 (Figure 1a). For MAYV mosquito challenges, we utilized two different strains of the virus (BeAn343102, genotype -D, and BeAr505411, genotype -L). Results are reported as 1) infection rate (IR), the proportion of challenged mosquitoes with infected midguts, 2) dissemination rate (DR) and efficiency (DE), the proportion of infected (DR) or challenged (DE) mosquitoes with infected bodies, and 3) transmission rate (TR) and efficiency (TE), the proportion of infected (TR) or challenged (TE) mosquitoes with infected saliva. These ratios were calculated and analyzed using viral titers measured from the midgut, carcass (rest of the body), and saliva, respectively, and are reported with numerical subscripts to indicate sampling day. Viral titers are described as means at individual timepoints (shown as M_x_, C_x_, or S_x_(_in_), where ‘M’ represents midgut, *‘C’* represents carcass, *‘S’* represents saliva samples, ‘×’ denotes the day of collection (days post infection), and ‘in’ represents only infected subsets. Average titers across all timepoints are reported without a timepoint subscript.

For both MAYV genotypes we found that *An. albimanus* was highly susceptible to infection, dissemination, and transmission. MAYV-D successfully established infections in the midgut and disseminated to the rest of the body in nearly all mosquitoes at all surveyed timepoints (Figure 2a; IR: 97.6%, DR: 97.6%, DE: 95.3%). While infection prevalence was consistent across time, disseminated viral titers rose until 10 dpi (Figure 2a; C_7_ vs C_10_: *U*=250, p=0.034), when both infection (M_10_: 1.3×10^6^ffu/mL) and dissemination (C_10_: 3.5×10^7^ffu/mL) titers were highest. The infection and dissemination patterns of MAYV-L were similar to those of MAYV-D, with the virus infecting and disseminating through mosquitoes at high rates (Figure 2b; IR: 94.3%, DR: 98.8%, DE: 93.2%). As with MAYV-D, MAYV-L titers also peaked at 10 dpi both in midguts (M_10_: 7.6×10^6^ ffu/mL; M_7_ VS M_10_: *U*=125, p<0.0001) and carcasses (C_10_: 2.5×10^6^ ffu/mL; C_7_ vs C_10_: *U*=119, p<0.0001). Titers then decayed significantly from this peak (M_10_ vs M_14_: *U*=151, p=0.0004; C_10_ vs C_14_: *U*=191.5, p=0.007). Despite high prevalence at all timepoints, we observed a higher variation in infection and dissemination for MAYV-D compared to MAYV-L (i.e., M_14_: F=3.23, p=0.0026, C_14_: F=4.84, p<0.0001). Transmission trends differed slightly between the two genotypes. Unlike its infection and dissemination rates, which were high and steady, MAYV-D transmission efficiency (Figure 2a) increased over time (TE_7_: 20%, TE_10_: 32%, TE_14_: 48.4%), though viral titers detected in those infectious mosquitoes remained constant (S_7in_: 1.3×10^3^ ffu/mL, S_14in_: 1.25×10^3^ ffu/mL). Note that MAYV-D was not sampled at 21 dpi due to high mortality that differed from controls (Figure 2c and Figure S1; 96.9%, χ^2^=43.72, df=1, p<0.0001). For MAYV-L, virus was present in the saliva samples of about one quarter of mosquitoes at most time points (TE_7_: 26.9%, TE_10_: 23%, TE_14_: 22.2%) but neither prevalence nor viral titers (S_7in_: 1.76×10^2^ ffu/mL, S_14in_: 3.11×10^2^ ffu/mL) increased significantly over time. At 21dpi, no mosquitoes were able to transmit the virus (i.e., TE_21_: 0%), which can be due to viral clearance from the body. Indeed, lower titers were also detected in midgut and carcass at 21 dpi (Figure 2b; MAYV-L: M_14_ vs M_21_: U=19.5, p<0.0001; C_14_ vs C_21_: *U*=8.5, p<0.0001). However, these low titers may reflect selection bias, as mortality was also very high in MAYV-L challenged mosquitoes at 21 dpi (Figure 2c and Figure S1; 93.75%, χ^2^=15.59, df=1, p<0.0001).

**Figure 2.**
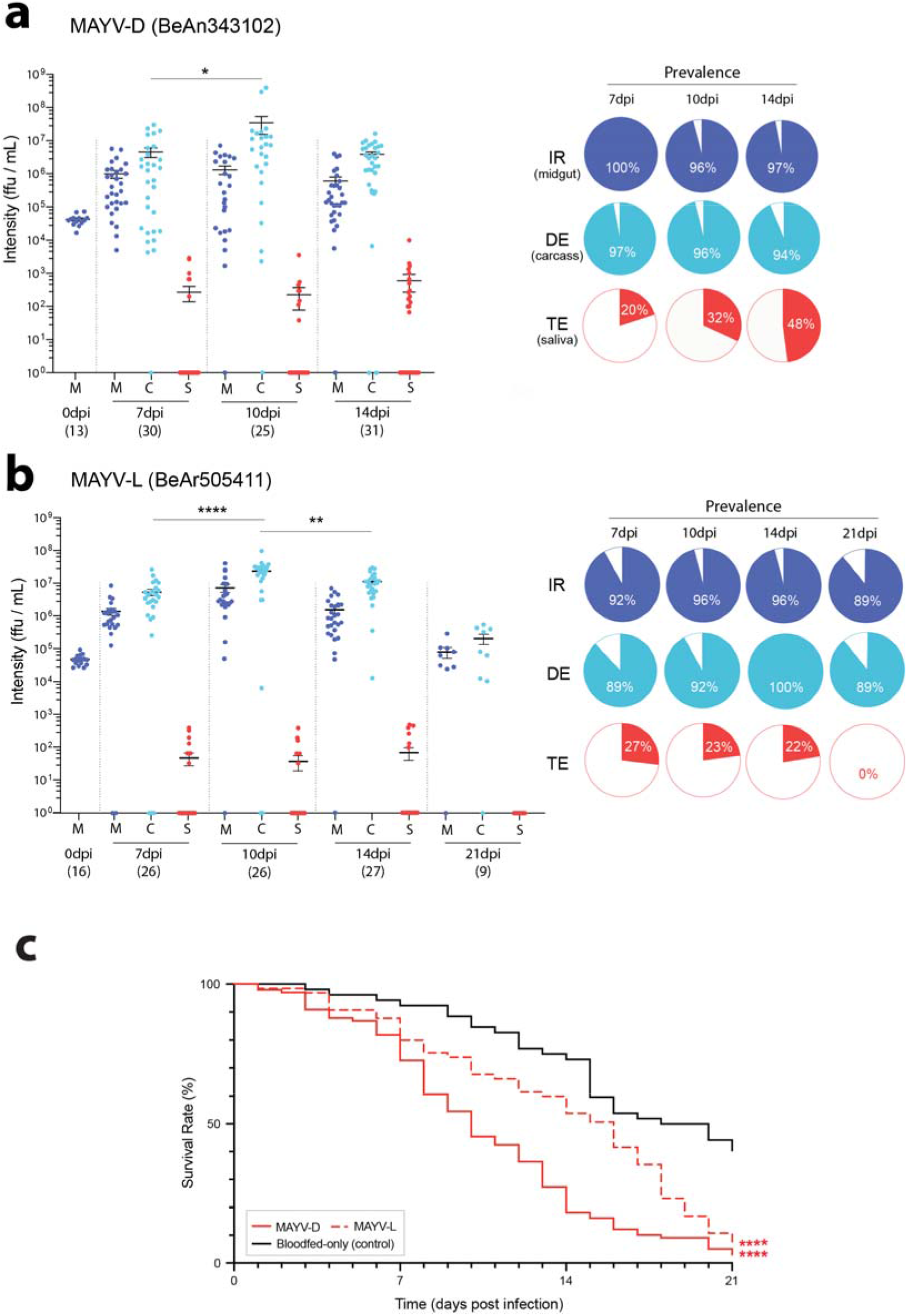
Viral titer in *An. albimanus’* midgut (M), rest of the body (carcass, C) and saliva (S) after exposure to **a)** MAYV-D, or **b)** MAYV-L. Each dot corresponds to the titer of a single mosquito sample, with number of collected samples (n) depicted below each timepoint. Pie charts indicate prevalence of infection (IR, dark blue), dissemination (DE, light blue) and transmission (TE, red) among the total challenged mosquitoes, **c)** *An. albimanus’* mortality associated to challenge and infection with MAYV strains. Statistical significance between virus-treated and bloodfed-only samples is indicated by stars (**** p<0.0001) and performed by curve comparison using a survival log-rank Mantel-Cox test.

### *An. albimanus* is not a competent vector of CHIKV or DENV2

While MAYV was able to successfully infect and transmit through *An. albimanus* mosquitoes, this was not the case for the other assayed viruses endemic in its native range. Following challenge with CHIKV, *An. albimanus* was able to become infected at moderate levels; we detected viral presence in 33% of the mosquitoes’ midguts 7 dpi (Figure 3a; My_7in_: 2×10^4^ffu/mL), but that decreased to 10% at 10 and 14 dpi (M_10in_: 3.67×10^4^ffu/mL and M_14in_: 1.44×10^4^ffu/mL). We found that CHIKV did not efficiently escape the midgut and disseminate, likely due to a midgut escape barrier. Only two carcass samples were CHIKV positive at 7dpi (6.7%, 2/30), and 0% of infections were disseminated at both 10 and 14 dpi (Figure 3a). None of these carcass-infected mosquitoes progressed to infected saliva. It therefore appears that CHIKV infection, at least with the H20235 virus strain and *An. albimanus* colony strain we tested, can only be established at the midgut level and rarely disseminates. CHIKV-associated mosquito mortality was not significantly different compared to the bloodfed-only controls (Figure 3c and Figure S1; 45.9%, χ^2^=0.82, df=1, p=0.365).

**Figure 3.**
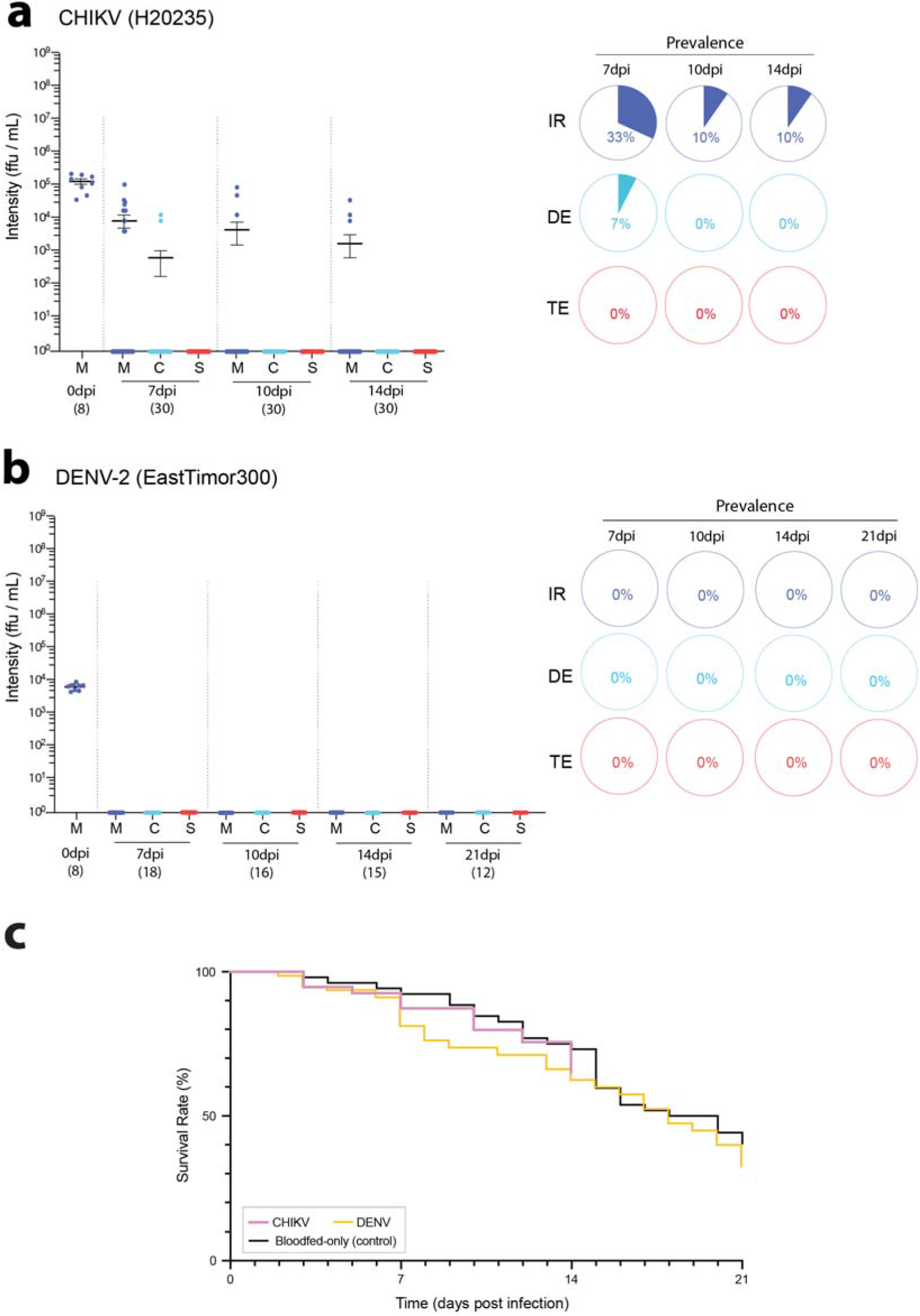
Viral titer in *An. albimanus’* midgut (M), rest of the body (carcass, C) and saliva (S) after exposure to **a)** CHIKV or **b)** DENV-2. Each dot corresponds to the titer of a single mosquito sample, with number of collected samples (n) depicted under each timepoint. Pie charts indicate prevalence of infection (IR, dark blue), dissemination (DE, light blue) and transmission (TE, red) among the total challenged mosquitoes, **c)** *An. albimanus’* mortality associated to challenge and infection with CHIKV and DENV-2. Statistical significance between virus-treated and bloodfed-only samples was performed by curve comparison using a survival log-rank Mantel-Cox test.

Besides American alphaviruses, we also assessed *An. albimanus’* vector competence for DENV2 due to its socioeconomic importance worldwide and because it is endemic in *An. albimanus’* habitat. As with other anophelines, *An. albimanus* was refractory to DENV infection. Despite challenging the mosquitoes with relatively high titers (5×10^6^ ffu/mL) and detecting positive sampling at 0dpi (which depicts mosquito intake of an infectious bloodmeal, Figure 3b), none of the collected individuals (Figure 3b; 0/61) from 7 to 21dpi was found to carry infectious virus in midgut, carcass, or saliva. These results led us to question whether anophelines presented a flavivirus-specific midgut barrier, which would restrict viral replication in the midgut upon the ingestion of an infective bloodmeal, or instead if they possess a body-wide infection or replication barrier against flaviviruses that would prevent replication in all tissues. To bypass the hypothetical midgut barrier, mosquitoes belonging to four *Anopheles* species *(An. gambiae, An. stephensi, An. guadrimaculatus* and *An. albimanus)* were injected with infectious viral stocks of the flaviviruses DENV or Zika (ZIKV) into their hemolymph. After 3 days of infection, none of the mosquitoes presented infectious particles in their bodies as assessed by FFA (Figure S2), indicating that the injected virus was not able to replicate within the mosquito and that the tested anophelines are completely refractory to flavivirus infections.

### Alphaviruses present outside the Americas can infect, disseminate, and be transmitted by *An. albimanus*

*We* also asked if *An. albimanus* may be a suitable vector of ONNV and SINV viruses. These alphaviruses are not known to have caused outbreaks in the Americas yet, but they could emerge in currently unaffected areas due to globalization, travel, and climate change—just as Zika virus spread to new continents^23,24^. We found infectious ONNV virions (p5’dsONNic) were able to both infect and disseminate from the midgut in nearly all challenged *An. albimanus* when they were fed at 1×10^7^ ffu/mL (Figure 4a; IR: 97.9%, DR: 98.9%, DE: 96.8%), showing that the species is susceptible to the virus. Unlike the pattern observed for MAYV, the highest ONNV infection intensity in the midgut was detected at 7dpi (Figure 4a; M_7_= 4×10^s^ffu/mL; M_7_ vs M_10_: *U*=90.5, p<0.0001), which then dropped slightly and remained stable throughout the remaining timepoints (M_10_ vs M_21_: *U*=167, p=0.29). Dissemination viral titers were similar to MAYV, peaking at 10dpi (C_10_= 2.5×10^6^ffu/mL; C_7_ vs C_10_: *U*=148.5, p=0.0004) followed by a stable plateau (C_10_ vs C_21_: *U*=155.5, p=0.18). Despite high viral prevalence in midgut and body, transmission rates were very low early in ONNV infections (TE/TR_7-14_: 4-7%) until an abrupt increase at 21dpi (TE/TR_21_: 31.3%; S_14_ vs S_21_: *U*=157, p=0.014). Our data thus shows that the virus can efficiently invade the salivary glands only late in the course of infections. Mortality was not significantly different in ONNV-positive mosquitoes compared to their bloodfed-only counterparts (Figure 4c and Figure S1; 70.4%, χ^2^=2.06, df=1, p=0.151).

**Figure 4.**
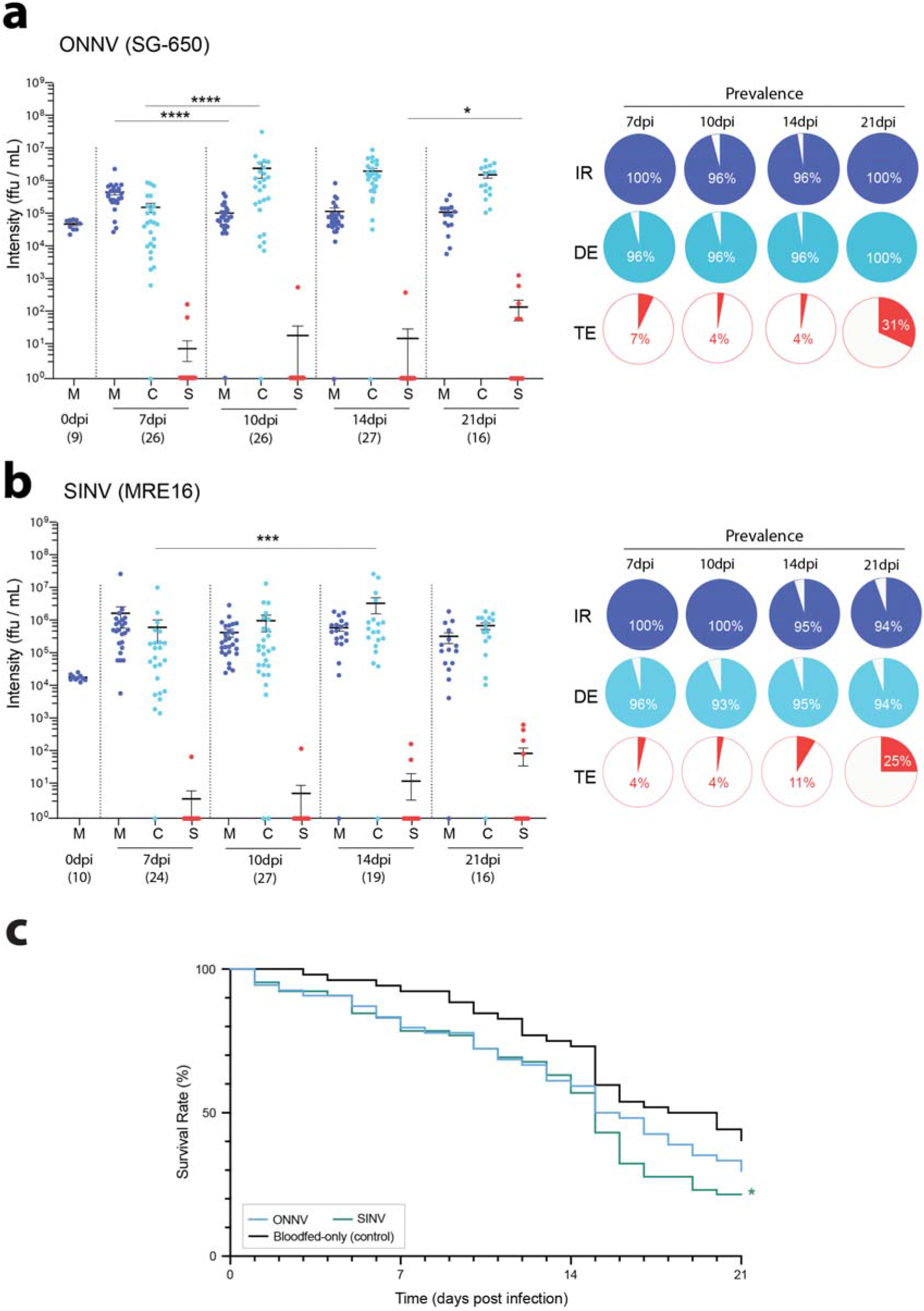
Viral titer in *An. albimanus^1^* midgut (M), rest of the body (carcass, C) and saliva (S) after exposure to **a)** ONNV or **b)** SINV. Each dot corresponds to the titer of a single mosquito sample, with number of collected samples (n) depicted under each timepoint. Pie charts indicate prevalence of infection (IR, light blue), dissemination (DE, dark blue) and transmission (TE, red) among the total challenged mosquitoes, **c)** *An. albimanus’* mortality associated to challenge and infection with MAYV strains. Statistical significance between virus-treated and bloodfed-only samples is indicated by stars (*p<0.05) and performed by curve comparison using a survival log-rank Mantel-Cox test.

*Anopheles albimanus* was also susceptible to be infected and transmit SINV, and infection dynamics were broadly similar to those of ONNV. Almost all mosquitoes presented infected midguts (Figure 4b; IR: 97.7%), with dissemination to the rest of their bodies also occurring at high frequencies (DR: 96.4%; DE: 94.2%). SINV titers were stable over time—only increasing in carcasses (i.e., dissemination) from 7 to 14dpi (*U*=115, p=0.005). This may indicate a slower replication rate within the mosquito tissues that delays and reduces its vectorial capacity. Early in the course of infections, few saliva samples presented infectious SINV (S_7-10_: 4%; S_14_: 11%) and those that did had low titers. Similar to ONNV, viral loads in saliva rose significantly at the latest timepoint (TE_21_: 25%, S_21in_: 3.7×10^2^pfu/mL), again representing a slower transition from midguts to the salivary glands.

## Discussion

Here we demonstrate that *An. albimanus,* the most common *Anopheles* mosquito in Mesoamerica and the Caribbean^6,25,26^, is a competent laboratory vector for a broad range of alphaviruses but refractory to flavivirus infection. Though arboviral spread through anophelines has received little research attention, we found this species was readily infected and transmitted three of five tested viruses—suggesting that it (and possibly other anophelines) may be susceptible to infection with a substantial number of viruses^3^. Though the tested alphaviruses (except CHIKV) were able to establish disseminated infections and be transmitted by this species, they presented different patterns of infection and transmission.

Notably, an alphavirus that is endemic to the Americas (MAYV)—and which therefore has the potential to infect *An. albimanus* in the wild—had the highest transmission efficiency of the tested viruses. Both MAYV genotypes (-D and -L; Figure 2a and 2b, respectively) are able to escape the mosquito’s tissue barriers^27^, replicating freely within the body of the mosquito and reaching the salivary gland lumen. Saliva samples tested positive for MAYV at a substantial rate (20-48%) in all surveyed timepoints except 21dpi, when most mosquitoes that potentially carried high viremia may have suffered from MAYV-associated mortality (Figure 2c and Figure S1). We also found MAYV infections progressed the most rapidly, as indicated by higher titers in midguts and body, as well as earlier and greater prevalence in saliva. However, it remains unclear whether (and how often) *An. albimanus* becomes infected with MAYV in the wild, or whether this pathogen may have been shaped by co-evolution with *An. albimanus* over time.

Our data also suggest there may be differences in how the two MAYV subgroups infect the tested colony of An. *albimanus.* For example, we detected higher variance in MAYV-D infection titers compared to those of MAYV-L infections (Figure 2a vs 2b). While both show a short extrinsic incubation period (EIP)^28^ that represents heightened vectorial capacity, MAYV-D’s transmission increased over time while MAYV-L remained consistent at around 25%, though this pattern was not statistically significant. Similar variation between MAYV genotypes in infection was previously observed in *An. gambiae*^20^. These differences suggest there may be interesting phenotypical variation in infection that could be explored in future studies.

To date, few MAYV outbreaks have been detected in humans, with most infections occurring in the Amazon basin^29^. However, recent studies indicate that MAYV true incidence may be underestimated^30^ because its symptomatology resembles that of other pathogens in the area (DENV, CHIKV). Epidemiological concern is rising due to MAYV’s ability to colonize urban areas and its potential to be transmitted through lesser-known vectors of arboviruses in Central and South America^29,31^. With the geographical range expansion of An. *albimanus* deeper into South America as well as northward into the USA, coupled with the high viral susceptibility and transmission observed in a laboratory colony, there is vectorial potential for this species to drive such outbreaks.

Not all alphaviruses endemic to the Americas were successful in infecting *An. albimanus.* Most notably, CHIKV had poor prevalence in challenged mosquitoes. Differences in transmission capacity can be explained by dynamic and complex virus-vector interactions that affect the vectorial capacity of the mosquito species^32^, and which may trace back to variation in host-to-pathogen genotype interactions (GxG). GxG interactions govern outcomes in many systems, including arboviral infections^33^ and symbiosis^34^ in insects. In some cases^26,27^ the combination of a specific insect genotype and a specific pathogen genotype strongly shapes infection status and progression. While *Anopheles* are not typically vectors for CHIKV^35^, some species including *An. stephensi* can harbor and transmit it^21^. In these rare cases, GxG interactions may be key in the ability of the mosquito to transmit the virus. Following what has been observed for most anophelines, *An. albimanus* were able to harbor the infection following a viral bloodmeal but were refractory to dissemination and transmission of CHIKV (Figure 3a). One hypothesis is that *An. albimanus* has co-evolved with this strain of CHIKV (collected on the Caribbean island of St. Martin) and the GxG interaction between mosquito and virus is not ideal^33,36^. Of course, our findings do not exclude the possibility that CHIKV may be transmitted through *An. albimanus* mosquitoes in some circumstances, as these results were obtained using a single laboratory colony. It remains possible that wild strains from other locations may present a more suitable combination for the virus to transmit—either by differing in their nuclear genotype^33^ or microbiome composition^37^.

We found that *An. albimanus* can also carry and transmit the non-American alphaviruses ONNV and SINV (Figure 4a-b). These two viruses affect different geographical and climate regions, and are primarily transmitted by other mosquito species^38,39^. ONNV is closely related to CHIKV and MAYV (all belong to the Semliki Forest antigenic complex) and causes outbreaks in humans in Central Africa. In contrast, SINV is a distant relative. It belongs to the Western equine encephalitis complex and is the causal agent of Pogosta disease, mostly in North Europe^40^ and South Africa^41^, with sporadic cases detected in Australia^42^. Despite their biological differences, both ONNV and SINV are able to infect *An. albimanus* albeit with slower transmission kinetics compared to MAYV. More specifically, though infected early at the midgut level, very few ONNV-or SINV-challenged mosquitoes were saliva-positive at the earliest timepoint tested (7dpi). Rather, their highest transmission was detected at 21dpi for both viruses. While neither ONNV nor SINV have been detected in the Americas yet, globalization and human travel^43^ pose a risk for the virus to spread there, especially SINV, whose primary vectors in Africa are the anopheline species An. *gambiae* and *An.funestus*^44,45^.

We found multiple anopheline species were refractory to flavivirus (DENV and ZIKV) infection, even when virus was injected directly into the hemolymph. This inability to harbor flaviviruses could trace back to a variety of different mechanisms, e.g., lack of a particular replication machinery component^46^ or absence of a specific cellular receptor or factor required for infection success^47, 48, 49^. However, there are reports of flavivirus infections in anophelines^3^, raising questions about the underlying mechanisms, and why some specific anopheline-flavivirus infections do occur.

In short, this study tested the capacity of *An. albimanus* to be an important vector of alphaviruses in the Americas. Our data show that, while *An. albimanus* is unlikely to drive CHIKV and flavivirus infections, other alphaviruses (especially MAYV) can infect and be transmitted through this mosquito species very efficiently. Our results highlight both the importance for *Anopheles* mosquitoes to be recognized as potential vectors of arboviruses, as well as the knowledge gaps that still need to be filled on the genus’ vectorial capacities worldwide.

## Materials and Methods

### Mosquitoes

The *Anopheles albimanus* mosquito colony (STECLA strain, MRA-126) was kept and reared at the Millennium Science Complex insectary (Center for Infectious Disease Dynamics, The Pennsylvania State University, PA, USA) at a continuous 27±1°C, 85% relative humidity, 12:12h light:dark cycle. Larvae were fed koi pellets (Tetra Pond Koi Vibrance; Tetra, Melle, Germany) from hatching to pupation. Adult mosquitoes were reared in 30×30×30cm metal cages and provided 10% sucrose solution *ad libitum,* as well as fed weekly with anonymous human blood (Biological Specialty Corporation, PA, USA) for reproduction and colony maintenance using a membrane feeder.

### Cells

African green monkey kidney cells (Vero; CCL-81) and *Aedes albopictus* larval cells (C6/36; CCL-126) (ATCC, VA, USA) were cultured in complete media consisting of DMEM or RPMI, respectively, complemented with 10% fetal bovine serum and 1% penicillin/streptomycin. For each passage, cells were detached by trypsinization (0.25 trypsin; Corning Inc., NY, USA) and diluted in fresh complete media or plated for experiments.

### Viruses

A total of six different viruses were used for experimental infections. Two strains of MAYV; BeAn343102 (BEI Resources, VA, USA) is a genotype D strain (MAYV-D) isolated in May 1978 from a monkey in Para, Brazil and BeAr505411 (BEI Resources, VA, USA) is a genotype L strain (MAYV-L) also isolated in Para, Brazil in March 1991 from *Haemagogus janthinomys* mosquitoes. The full-length ONNV and SINV infectious clones (p5’dsONNic and p5’dsMRE16ic) derive from the Uganda SG-650 strain of ONNV and wild-type MRE16 strain of SINV isolated from *An. malayensis*^50^, respectively. Both infectious clones were obtained on filter paper, transfected, and virions collected prior to passaging in Vero cells for the experiments. The DENV serotype 2 ET300 strain was isolated from a human patient in Timor-Leste in 2000 (GenBank accession: EF440433.1). Lastly, the CHIKV H20235 strain (NR-49901; BEI Resources, Manassas, VA, USA) was isolated from a human in St. Martin in 2013. All work with this strain (from cell culture to mosquito infections) was performed in the Eva J Pell Laboratory for Advanced Biological Research BSL-3 facility at The Pennsylvania State University.

All alphaviruses were passed in African green monkey kidney (Vero) cells at 37°C in a humidified 5% CO_2_ incubator, whereas DENV was passed in *Ae. albopictus* RNAi-deficient C6/36 cells at 28°C. Viruses were allowed to infect cells at a MOI of 0.1 for 1h and then removed and replaced with media containing 2% FBS. Virus-infected supernatant was aliquoted at different timepoints (typically 24h post-infection for alphaviruses and 7 days post-infection for DENV) and stored at −80°C until further titration or use for mosquito infections. Viral stock titers were obtained using focus forming assays (ffu/mL) or plaque assays (pfu/mL), described below.

### Vector competence assays

To determine the vectorial capacity of *An. albimanus,* adult females were orally challenged with an infected bloodmeal containing a high titer dose of one of the five togaviruses or DENV. Specifically, six to eight day-old non-blood fed females were allowed to feed on human blood for 1h through a synthetic membrane at the bottom of a glass feeder jacketed with 37°C water, and containing either 1×10^7^ ffu/mL (alphaviruses) or 5×10^6^ ffu/mL (DENV) of the stocks obtained above (Figure 1c, grey boxes). Fully engorged mosquitoes were sorted from non-fed ones and split evenly in cups for each collection timepoint. A small subset of mosquitoes were collected at 0dpi to confirm that the viral intake was infectious and similar across samples.

Infection (IR), dissemination and transmission rates (DR and TR), as well as dissemination and transmission efficiency (DE and TE), were assessed at 7, 10, 14 and 21 dpi. IR was measured as the rate of mosquitoes with infected midguts among the total number of mosquitoes. DR and DE were measured as the rate of mosquitoes with infected carcasses among the mosquitoes with infected midguts, or over the total assessed samples, respectively. TR and TE were measured as the rate of mosquitoes with infectious saliva among the positive bodies or the total number of assessed mosquitoes, respectively.

At all timepoints, mosquitoes were anesthetized using triethylamine (Sigma, St. Louis, MO, USA) before individual forced salivation. Saliva was collected by placing the female’s proboscis into a pipette tip containing 20μL of a 50% sucrose 50% FBS solution, as previously described^51^, for 30min. Solution was then released into a tube filled with 100μL of mosquito diluent (20% heat-inactivated fetal bovine serum, 50 μg/mL penicillin/streptomycin, 50 μg/mL gentamicin, and 2.5 μg/mL fungizone in Dulbecco’s phosphate-buffered saline) and placed on ice. Each female’s midgut was dissected and placed in a 2mL tube containing 300μL of mosquito diluent. The rest of the body (carcass) was also collected on an identical tube. Tissue samples were homogenized at 30Hz for 2min using a TissueLyser II (QIAGEN, Hilden, Germany). All samples were stored at −80°C until viral testing.

### Intrathoracic injections

*An. gambiae, An. stephensi, An. guadrimaculatus* and *An. albimanus* females were briefly anesthetized on a chill block (BioQuip Products, CA, USA) cooled to 4°C and DENV and ZIKV stocks were injected intrathoracically under a microscope using a pulled glass capillary with a manual microinjector (Nanoject II, Drummond Sci., PA, USA) to ensure uniformity of dosage. Sixty-nine microliters of diluted virus stock (~70 DENV pfu) were delivered intrathoracically into each female. After injection, mosquitoes were maintained under standard housing conditions of 27°C with 80% relative humidity, 12 h light/dark cycle and fed 10% sucrose solution *ad libitum*.

### Focus forming assay

Presence of infectious particles of all viruses except SINV in saliva, midguts and carcasses was tested by focus forming assays in Vero (alphaviruses) or C6/36 (DENV) cells. Cells were counted using a hemacytometer (Hausser Scientific, PA, USA) and plated in complete media the day before infection to achieve 80-90% confluency (Vero: 3×10^4^ cells/well, C6/36: 2×10^s^ cells/well) in 96-well plates. The following day, media was removed from all wells and cells were incubated for 1h with 30μL of 10-fold dilutions (10^-1^ to 10^-4^) of each homogenized tissue sample in FBS-free media. Saliva samples were not diluted due to their lower titers. Viral media was removed from the wells after 1h, replaced with 100μL of overlay (final 0.8% methylcellulose (or CMC) in complete media) and incubated at 37°C for 24h or 28°C for 3 days, depending on the cell culture used. Cells were then fixed using 4% paraformaldehyde in PBS (Sigma-Aldrich, MO, USA) for 20min and permeabilized with 0.2% Triton-X in PBS for another 20min. Samples were washed 2-3 times with cold 1x PBS after both fixation and permeabilization steps. Viral antigens in infected cells were labeled overnight using mouse monoclonal anti-Chikungunya virus E2 envelope glycoprotein clone CHK-48 (for all alphaviruses except SINV; α-CHK-48, BEI Resources, VA, USA) or mouse monoclonal anti-flavivirus clone D1-4G2-4-15 (for DENV; BEI Resources, Manassas, VA, USA) diluted 1:500 in PBS. The next day, cells were again washed thoroughly with cold PBS to remove unbound primary antibody. Bound primary antibody was then labeled for 1h at room temperature using Alexa488 goat anti-mouse IgG secondary antibody (Invitrogen, OR, USA) at a 1:750 dilution in PBS, which was then rinsed off with RO water before evaluation by fluorescence microscopy. Green fluorescence was observed using a FITC filter on an Olympus BX41 microscope with a UPlanFI 4x objective. Foci were counted by eye in the appropriate dilution (10-100 foci) and viral titers back calculated to focus forming units per mL (ffu/mL).

### Plaque assay

The α-CHK-48 antibody used in FFAs does not cross-react with SINV, which is evolutionarily the most distantly related to CHIKV^52^. Thus, we elected to assess mosquito SINV infections by traditional plaque-forming assays (PFA), which are comparable to FFA since both detect presence of infectious viral particles in a sample using a cell-based method.

Mosquito samples were tested for SINV infectious particles by plaque assay on Vero cells with minimal modifications^53^. The day before infection, cells were counted as described above and plated (5×10^6^ cells/well) in 6-well plates. For saliva infections, media was removed and replaced for 100μL of undiluted sample. For midguts or carcasses, 10-fold dilutions using FBS-free media were performed and 180μL of each dilution (in most instances: 10^-2^ to 10^-4^) was used for cellular infection. Inoculated plates were placed in a 37°C incubator with 5% CO_2_ for 1h for viral entry to occur. Then, virus-containing media was removed and replaced with 1.5mL of an agar overlay (equal parts of complete media and 1.2% agarose) and placed back into the incubator. After two days, 1.5mL of a second agar overlay (identical to the first agar overlay but containing 1% final concentration of neutral red (Amresco Inc., OH, USA) to allow for cellular staining), was added to each well and plates incubated. The following day, agar discs were removed, and samples treated with 4% formaldehyde to inactivate any remaining virus. Each stained well was rinsed thoroughly with water and set aside to dry well. Wells that produced 10-100 plaques were used to ensure accurate counts, and viral titers for mosquito saliva, midguts and carcasses were calculated in plaque-forming units per mL (pfu/mL). When samples produced too many plaques to count, additional plaque assays were performed with extra 10-fold dilutions.

### Statistical analyses and figure generation

Differences in viral titer between midgut, carcass and saliva samples were assessed by twotailed non-parametric Mann-Whitney *U* tests due to the non-normality of the samples. Survival data for each virus was compared pairwise to a bloodfed-only control sample using a log-rank Mantel-Cox test. All p-values that were below 0.05 (p<0.05) were considered significant. All data was initially plotted and analyzed using Prism software version 9.2.0 (283) (GraphPad Inc., CA, USA). Final figures were assembled using Adobe Illustrator 2021 (25.4.1, Adobe Inc., CA, USA).

**Table 1.** Parameters describing infections in *Anopheles albimanus.* Infection rate (IR), dissemination rate (DR), dissemination efficiency (DE), transmission (TR) rate, and transmission efficiency (TE) for all the assessed viruses are reported. N = total number of challenged mosquitoes, x = number with virus present in midgut, y = number with virus present in carcass, z = number with virus present in saliva, dpi = days post infection.

|  | % IR (x/N) | % DR (y/x) | % DE (y/N) | % TR (z/y) | % TE (z/N) |
| --- | --- | --- | --- | --- | --- |
| **MAYV-D** |  |  |  |  |  |
| 7dpi | 100% (30/30) | 96.7% (29/30) | 96.7% (29/30) | 20.6% (6/29) | 20% (6/30) |
| 10dpi | 96% (24/25) | 100% (24/24) | 96% (24/25) | 33.3% (8/24) | 32% (8/25) |
| 14dpi | 96.8% (30/31) | 96.7% (29/30) | 93.5% (29/31) | 51.8% (15/29) | 48.4% (15/31) |
| 21dpi | - | - | - | - | - |
| **MAYV-L** |  |  |  |  |  |
| 7dpi | 92.3% (24/26) | 95.8% (23/24) | 88.5% (23/26) | 30.4% (7/23) | 26.9% (7/26) |
| 10dpi | 96.2% (25/26) | 96% (24/25) | 92.3% (24/26) | 25% (6/24) | 23% (6/26) |
| 14dpi | 96.3% (26/27) | 104% (27/26) | 100% (27/27) | 22.2% (6/27) | 22.2% (6/27) |
| 21dpi | 88.9% (8/9) | 100% (8/8) | 88.9% (8/9) | 0% (0/8) | 0% (0/9) |
| **ONNV** |  |  |  |  |  |
| 7dpi | 100% (26/26) | 96.2% (25/26) | 96.2% (25/26) | 8% (2/25) | 7.7% (2/26) |
| 10dpi | 96.2% (25/26) | 100% (25/25) | 96.2% (25/26) | 4% (1/25) | 3.8% (1/26) |
| 14dpi | 96.3% (26/27) | 100% (26/26) | 96.3% (26/27) | 3.8% (1/26) | 3.7% (1/27) |
| 21dpi | 100% (16/16) | 100% (16/16) | 100% (16/16) | 31.3% (5/16) | 31.3% (5/16) |
| **SINV** |  |  |  |  |  |
| 7dpi | 100% (24/24) | 95.8% (23/24) | 95.8% (23/24) | 4.3% (1/23) | 4.2% (1/24) |
| 10dpi | 100% (27/27) | 92.6% (25/27) | 92.6% (25/27) | 4% (1/25) | 3.7% (1/27) |
| 14dpi | 94.7% (18/19) | 100% (18/18) | 94.7% (18/19) | 11.1% (2/18) | 10.5% (2/19) |
| 21dpi | 93.8% (15/16) | 100% (15/15) | 93.8% (15/16) | 26.7% (4/15) | 25% (4/16) |
| **CHIKV** |  |  |  |  |  |
| 7dpi | 33% (10/30) | 20% (2/10) | 6.7% (2/30) | 0% (0/2) | 0% (0/30) |
| 10dpi | 10% (3/30) | 0% (0/3) | 0% (0/30) | - | 0% (0/30) |
| 14dpi | 10% (3/30) | 0% (0/3) | 0% (0/30) | - | 0% (0/30) |
| 21dpi | - | - | - | - | - |
| **DENV** |  |  |  |  |  |
| 7dpi | 0% (0/17) | - | 0% (0/17) | - | 0% (0/17) |
| 10dpi | 0% (0/14) | - | 0% (0/14) | - | 0% (0/14) |
| 14dpi | 0% (0/13) | - | 0% (0/13) | - | 0% (0/13) |
| 21dpi | 0% (0/8) | - | 0% (0/8) | - | 0% (0/8) |

## Acknowledgements

ONNV and SINV infectious clones were kindly provided by Brian Foy (Colorado State University, Ft. Collins, CO, USA). DENV-2 ET300 strain was provided by the McGraw Lab (Penn State University, University Park, PA, USA). Mosquitoes and antibodies were provided by BEI Resources. We would like to thank Kaylee Montanari and Amelia Romo from the Rasgon Lab, as well as personnel from the Eva J Pell BSL-3 laboratory for Advanced Biological Research at Penn State, for their continuous technical support.

## Author contributions

G.T. and J.L.R. conceptualized the study. G.T., M.N., and M.B. performed the experiments. G.T. and H.C.M. analyzed the data. G.T. and H.C.M. designed all artwork and figures for the paper. G.T., H.C.M. and J.L.R. contributed to the writing. All authors approved the final manuscript.

## Author’s current addresses

GT, MN, HCM and JLR: *Department of Entomology, the Center for Infectious Disease Dynamics, and the Huck Institutes of the Life Sciences, The Pennsylvania State University, University Park, Pennsylvania, USA* MB: Instituut voor Tropische Geneeskunde, Antwerp, Belgium

