## Supplementary material for "*Anopheles albimanus* is a potential alphavirus vector in the Americas": Sup fig 1 and 2

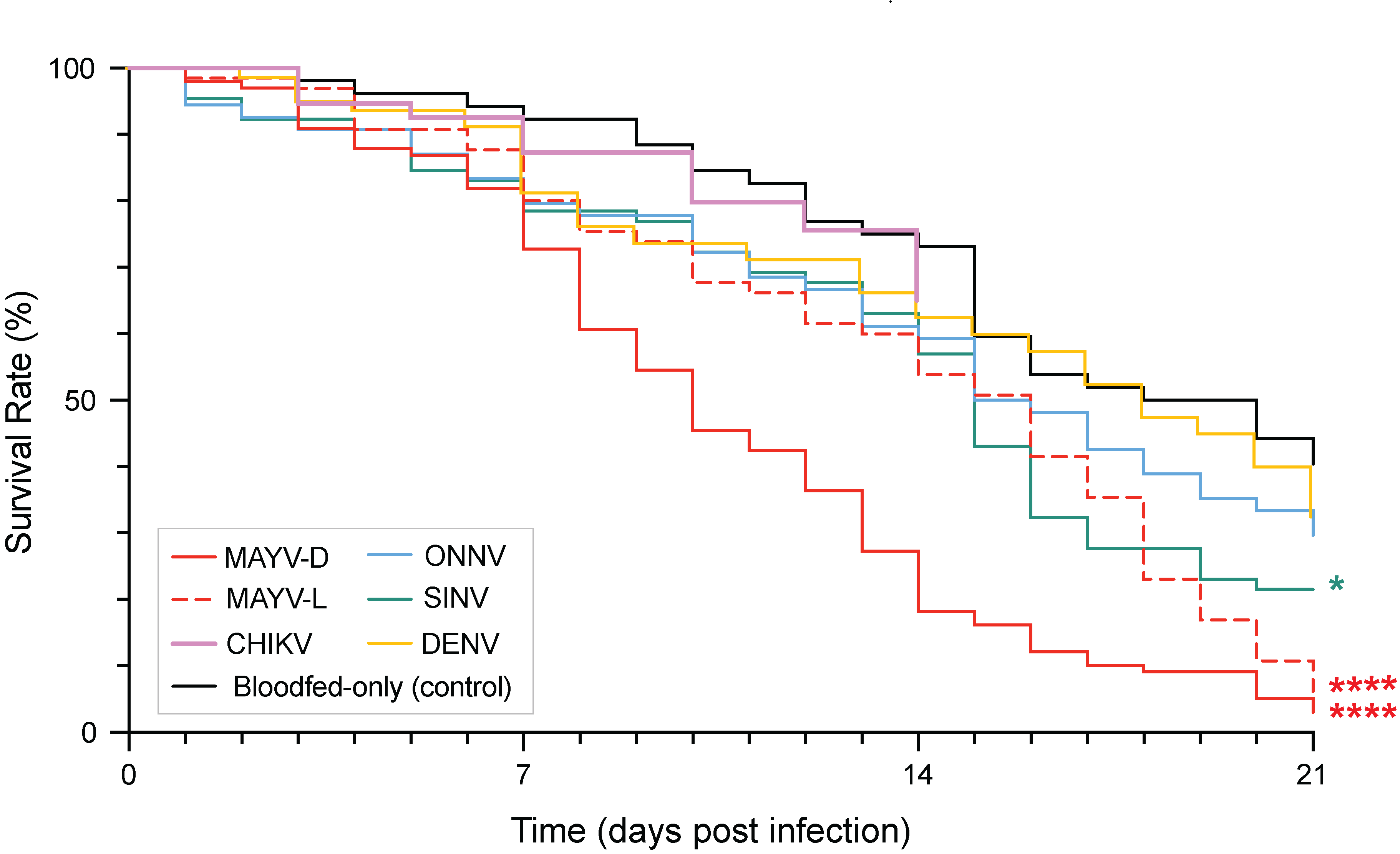

**Figure S1. –** *An. albimanus’* mortality associated to challenge and infection with the different viruses. Statistical significance between virus-treated and bloodfed-only samples is indicated by stars (**** p<0.0001; * p<0.05) and performed by curve comparison using a survival log-rank Mantel-Cox test.

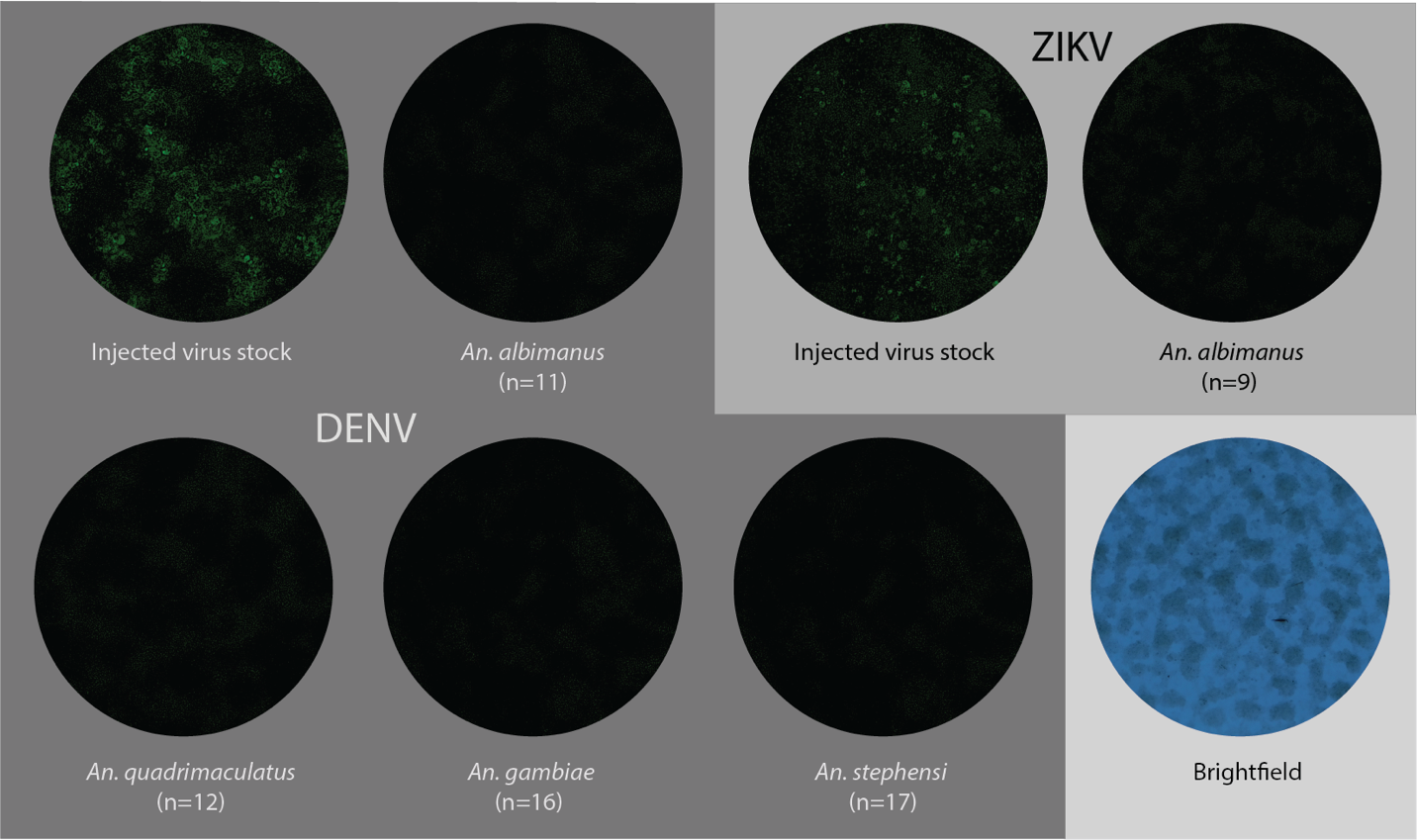

**Figure S2. –** Four diverse *Anopheles* species were injected intrathoracically with DENV and ZIKV to assess for presence of a flavivirus-specific midgut barrier in the genus. The figure depicts FFA on samples of intrathoracically-injected mosquitoes. FFAs on viral stocks serve as positive controls for both techniques and infective virus at time of injection, with each virus stained in green using a specific primary antibody coupled with an Alexa Fluor 488 secondary antibody.
